# Annotated reference genome for the Critically Endangered, paleoendemic Bermuda skink (*Plestiodon longirostris*)

**DOI:** 10.64898/2026.11.12.766285

**Authors:** Owen Greenwood, Keiazia Burchall-Busby, Haley Raynor, Kameyah Jones-Mussenden, Leah J. Williams, Richard A. Griffiths, Mark E. Outerbridge, Jean-Pierre Rouja, Helena S. Turner, Bradley Cain, Gerardo Garcia, Miten Jain, Carika Weldon, Sarah M. Griffiths

## Abstract

Skinks, one of the largest vertebrate families with over 1,500 described species, are represented by only 19 publicly available chromosome-level genome assemblies. Here, we present the first whole-genome assembly of the Bermuda skink*, Plestiodon longirostris,* a Critically Endangered, evolutionarily distinct, and endemic species. We conducted a *de novo* genome assembly using Oxford Nanopore Technology long reads, followed by reference-guided scaffolding against the five-lined skink *Plestiodon fasciatus* and Gilbert’s skink *Plestiodon gilberti* genomes. We generated a 1.496 Gb assembly with 2,579 scaffold contigs over 500 bp in length, a scaffold N50 of 224.1 Mb, with 98.7% Benchmarking Universal Single-Copy Orthologs (BUSCO) completeness score and 94.8% of gene annotations transferred from the *P. fasciatus* annotated genome. Repetitive elements comprised 48.37% of the genome, including 33.79% unclassified repeats, while 9% of transposable element sequences contained internal stop codons, consistent with other reptile genomes. Synteny analysis revealed strong conservation of macrochromosomes and gene order through the *Plestiodon* genus, and progressive chromosomal rearrangement with increased phylogenetic distance across the wider skink family. We also present the first mitogenome for the species. These genomic resources will further evolutionary research in the skink family and will facilitate applied conservation genomic approaches in this Critically Endangered species.

**Article summary:** This article provides the first annotated, chromosome-level genome assembly of the Bermuda skink (*Plestiodon longirostris*), an endemic, Critically Endangered skink species. This report provides reference autosomal and mitochondrial genomes for researchers studying the population genomics of the Bermuda skink, as well as other closely related species. The *de novo* genome was fragmented but was highly complete and annotated once scaffolded against the existing *Plestiodon fasciatus* genome. This provides a high quality, annotated reference genome for other researchers studying lizard or skink genetics.

## Introduction

Skinks (family Scincidae) are a large, diverse family of lizards, comprising over 146 genera and 1,500 species. Skinks have evolved in all continents except Antarctica, across almost all habitats (Chapple et al. 2023). Due to their diversity, skinks are models for studying the evolution of a range of traits, such as sex determination systems, reproductive modes and thermal adaptation (Hill et al. 2018; Cabrelli and Hughes 2015). However, chromosome-level genome assemblies are only available for 19 species (∼1.3% of the family) in the National Center for Biotechnology Information (NCBI) database, highlighting a lack of high-resolution data to resolve ongoing questions on systematics and phylogenetics in the family and impeding comparative genomics research. Additionally, over 20% of skink species are threatened with extinction (Chapple et al. 2021), underscoring the need for further genomic resources to be developed to aid applied conservation genomics research in this group.

The Bermuda skink (*Plestiodon longirostris,* Cope 1861) is a Critically Endangered species endemic to the subtropical North Atlantic archipelago of Bermuda (Cox and Wingate 2021). Threatened by habitat loss, predation by invasive species, and entrapment in discarded bottles, recent population estimates show that the species is in continued decline, with fewer than 300 individuals remaining in the largest population (Greenwood et al. In Press; Turner et al. 2021), necessitating urgent conservation action to avoid extinction in the wild. Habitat and invasive species management are ongoing, however, given the perilous state of the species, the more radical measures of translocating individuals between populations and reintroduction from captivity are also under active consideration (Garcia et al. 2024). Genetic information can be crucial for effective implementation of these methods; it can show patterns of dispersal and isolation across fragmented habitats, identify local adaptation, and reveal levels of genetic diversity and inbreeding (Baguette et al. 2013). However, the lack of genomic resources for the Bermuda skink impedes understanding of these factors, leading to uncertainties into how to most effectively undertake these conservation activities.

As well as being endemic and Critically Endangered, the Bermuda skink is the last remaining species in its lineage, having diverged between 11 - 20 mya (Brandley et al. 2010). This makes it a highly evolutionarily distinct species (Gumbs et al. 2024), and an example of paleoendemism, where representatives of ancient highly diverged lineages are now restricted to a small geographic area, having previously been more widespread. As the only member of its lineage, conserving the Bermuda skink is important for preserving millions of years of evolutionary diversity, but moreover, genomic resources for the species will aid evolutionary biology research in the Scincidae more generally.

Here, using Oxford Nanopore long-read sequencing, we generate the first chromosome-level, annotated genome for the Bermuda skink. We also present the first mitochondrial genome for the species, assembled from the same Oxford Nanopore long reads. These genomic resources will support future applied conservation genomics research in the species, as well as study of Scincidae evolution.

## Materials and methods

### DNA Extraction and Sequencing

#### Long-read sequencing

We obtained genetic material from an opportunistically sampled autotomized tail of a Bermuda skink (*Plestiodon longirostris*) found while carrying out Capture-Mark-Recapture surveys on Southampton Island, Bermuda (32° 20′ 32.00′, -64° 40′ 04.00′) in August 2023. The sample was preserved in Nucleic Acid Preservation Buffer (Camacho-Sanchez et al. 2013) and stored at -20 °C. This was used to generate long-read sequence data for genome assembly. The sex of the individual the tail tip was obtained from is unknown, as Bermuda skink external appearance is not sexually dimorphic, and sex can currently only be determined either by observed egg laying, or in a necropsy.

We extracted high molecular weight DNA from 10 ng of tail tissue using [ammonium acetate precipitation](https://support.microsoft.com/en-us/internet-explorer/this-website-doesn-t-work-in-internet-explorer). DNA concentration was measured using Qubit fluorometric quantitation (Thermo Fisher Scientific). A library was prepared with the Oxford Nanopore Technologies (ONT) Ligation DNA Sequencing Kit (SQK-LSK114) and sequenced on 1 PromethION flow cell (FLO-PRO114M) using the PromethION 24 platform.

#### Short-read sequencing

For polishing, we carried out low-coverage whole genome sequencing from 198 buccal swab samples originally collected from wild Bermuda skinks as part of a population genetics study (Greenwood et al., unpublished). DNA was extracted using the DNeasy Blood and Tissue Kit (QIAGEN, Valencia, CA) or [ammonium acetate precipitation](https://www.protocols.io/view/chloroform-free-dna-extraction-ammonium-acetate-pr-x54v9pekzg3e/v2), and libraries were prepared with the NEBNext Ultra II FS DNA library preparation kit (New England Biolabs, Hitchin, UK). Samples were sequenced on an Illumina NovaSeq X Plus (150 bp paired-end reads) across two lanes of a 25B flow cell at the Centre for Genomic Research, University of Liverpool.

All samples were collected under a Scientific Research permit issued by the Government of Bermuda Department of Environment and Natural Resources.

### Autosomal Genome Assembly

We used chopper (v0.12; De Coster C Radermakers, 2023) to trim the Nanopore long reads, filtering with a minimum average Phred quality score of 15 and minimum read length of 500 bp, and trimming 27 bases from the start of each read and 38 bases from the tail. We then *de novo* assembled the long-read data using flye (v2.9.6; Kolmogorov et al., 2019) setting genome size at 1,500 Mb, as seen in other *Plestiodon* species. We used QUAST (v5.3.0; Gurevich et al., 2013) to determine the number of contigs and N50, and estimated genome completeness using BUSCO (v6.0.0; Tegenfeldt et al., 2025). We polished our genome with medaka (v2.2.0, Oxford Nanopore Technologies, 2025) then compared our original assembly with the polished one to determine quality improvement using QUAST (v5.3.0; Gurevich et al., 2013).

We also assembled chopper-trimmed reads against the existing annotated genomes of the five-lined skink *Plestiodon fasciatus* (Hoffman et al., 2026) and Gilbert’s skink *Plestiodon gilberti rubricaudatus* (Richmond et al., 2026), two of the most closely related skink species to the Bermuda skink. We used RagTag (v2.1.0; Alonge et al., 2022) to scaffold our assembly against these genomes, before using minimap2 (v2.30; Li, 2018) to generate dotplots to assess the chromosomal fit.

#### Gene Annotation

The minimap2 dotplots showed that all 13 chromosomes of the scaffolded assembly each corresponded to a single chromosome in both *P. fasciatus* and *P. gilberti rubricaudatus.* We used Liftoff (v1.6.3; Shumate C Salzberg, 2021) to annotate protein-coding genes using both reference *Plestiodon* genomes separately for comparison. We used minimap2 (v2.30; Li, 2018) alignments to select best placement using sequence identity and coverage thresholds. We assessed gene transfer efficiency by comparing the number of genes in the reference genomes to the number of genes lifted to the Bermuda skink genome.

#### Repeat Element Annotation

We used RepeatModeler (v2.0.7; Smit C Hubley, 2015) with the LTRStruct flag to improve long terminal repeat (LTR) retrotransposon detection. We combined this de novo library with the Squamata database in Dfam (v3; Storer et al., 2021) to generate a combined repeat library. We used RepeatMasker (v4.2.2; Smit et al., 2015) to identify and soft mask repeat sections using the -xsmall flag, and summarized repeat content by class (Long Interspersed Nuclear Elements (LINE), Short Interspersed Nuclear Elements (SINE), LTR, DNA transposon and unknown), expressed as a percentage of the total genome size. We visualised the age of transposable elements (TEs) as a Kimura 2-parameter (K2P) divergence landscape. The divergence of a given TE from the consensus sequence acts as a proxy for insertion age, with lower divergence values indicating more recently active, and therefore younger, elements. We visualised repeat landscapes and genome composition in R (v4.4.3; R Core Team, 2025) using ggplot2 (v4.0.3; Wickham, 2016). We polished the scaffolded genome with pilon (1.24; Walker et al., 2014) using the paired-end short-read Illumina sequence data from 175 individuals, mapped with BWA-MEM (0.7.19; Li, 2013) to the assembly (--fix all, minimum depth 5×; reads subsampled to 8%). We then assessed assembly quality with BUSCO and QUAST, followed by another round of RepeatModeler and RepeatMasker based on our genome, before re-running Liftoff to ensure gene matching was as complete as possible. Finally, we screened the genome for contamination using NCBI FCS-GX (v0.5.5; Astashyn et al., 2024). We identified and excluded five small contigs (3.2-48.6 kb) as bacterial contamination and masked one 877 bp region on chromosome 3. Our cleaned assembly retained 2,574 scaffolds.

### Genomic Landscape

We calculated gene density and repeat content using 1 Mb sliding windows, with a 500 kb step across all 13 macrochromosomes using the data.table package (v1.18.99; Barrett et al., 2026) in R. We determined gene count in each window by counting GFF3 gene features which overlapped the window coordinates, and we calculated repeat content using the proportion of base pairs in the window annotated as repetitive by RepeatMasker. We avoided double counting by clipping to window boundaries. We plotted gene density (genes per Mb) and repeat content (%) as area plots, faceted by chromosome, using ggplot2.

### Synteny analysis

We carried out pairwise synteny analyses among the Bermuda skink and comparison species (Gilbert’s skink *P. gilberti,* five-lined skink *P. fasciatus,* common blue-tongued skink *Tiliqua scincoides,* Puerto Rican Skink *Spondylurus nitidus,* and Cape Cliff lizard *Hemicordylus capensis)* using two complementary approaches. First, we carried out whole-genome alignments using minimap2 for sequence-level synteny, using asm5 preset to compare between assemblies of the same genus. We output alignments in PAF and filtered to retain only chromosome-level alignments with a minimum mapping quality of 30 and minimum alignment length of 10 kb.

We used GENESPACE (v1.3.1; Lovell et al., 2022) to identify syntenic blocks across the six species. We extracted protein sequences from each annotated genome using gffread (v0.12.9; Pertea C Pertea, 2020). We used transcript-level identifies from mRNA features as protein FASTA headers for genomes with funannotate- or Liftoff-derived annotations. We extracted protein accessions by joining mRNA coordinates to CDS protein_id attributes from NCBI RefSeq annotated genomes (*T. scincoides* and *H. capensis*). We generated BED files with four columns (chromosome, start, end, gene ID), with the longest isoform entry used per gene. We used OrthoFinder (v2.5.5; Emms C Kelly, 2019) to infer orthogroup, using DIAMOND (v2.1.24; Buchfink et al., 2021) for all-versus-all protein searches and MCScanX (v1; Wang et al., 2012) for syntenic block detection. We visualized the synteny map as a riparian plot using GENESPACE’s plot_riparian function, using *P. longirostris* as the reference genome. We used ImageMagick (v6.9.10; ImageMagick Studio LLC) and R to correct background and label formatting. We added a time-calibrated phylogenetic tree to the figure using ggplot and cowplot (v1.2; Wilke 2025), using divergence times obtained from TimeTree (Kumar et al. 2022) and the fossil-calibrated estimate for Bermuda skink lineage divergence of 11.5-19.8 Ma (Brandley et al. 2010).

We also determined the Bermuda skink putative X chromosome block using whole-genome pairwise synteny against the eastern three-lined skink *Bassiana duperreyi* genome, which has hypothetical sex chromosomes identified (Dissanayake et al. 2020; Hanrahan et al. 2025). We used minimap2 (v2.30; Li, 2018) with the asm20 preset, appropriate for the ∼150 Ma divergence between the Bermuda skink and *B. duperreyi* (Kumar et al. 2022). We identified the best matching *B. duperreyi* sequence for each Bermuda skink macrochromosome by extracting the highest scoring alignment per query chromosome from the PAF output.

### Mitochondrial Genome

We assembled the mitochondrial genome of the reference individual from the Oxford Nanopore reads. Reads were mapped with minimap2 (2.30; Li, 2018) against the Bluetail Mole skink *Plestiodon egregius lividus* mitogenome (GenBank Accession No.: AB016606.1), and a haploid consensus was called with bcftools (1.23.1; Danecek et al., 2021). We polished the consensus with Medaka (2.2.1; Oxford Nanopore Technologies, 2026) using the r1041_e82_400bps_hac_v4.3.0 model. To validate the sequence, we mapped the mitochondrial reads back to the polished consensus and called variants. Where the majority of reads (>50%) supported a different base at a position with at least 50× depth, we corrected the consensus to the majority base. We annotated the final mitogenome using MITOS2 (v2.1; Donath et al., 2019) with the vertebrate mitochondrial genetic code (code 2) and the RefSeq63 metazoan reference database.

## Results and Discussion

### Autosomal Genome

We generated 30.4 million Oxford Nanopore Technology (ONT) reads, totaling 67.56 Gb (mean read length 2,225 bp), at 45x coverage. The *de novo* assembly was 1.496 Gb, highly fragmented with 13,098 contigs, with a GC content of 45.68%, an N50 of 334 kb, and a BUSCO completeness of 94.1%. After scaffolding this assembly against the two existing *Plestiodon* reference genomes, we achieved 2,579 and 3,391 scaffold contigs, scaffold N50s of 176 Mb and 224 Mb, and BUSCO completeness scores of 98.2% and 98.7% for *P. gilberti* and *P. fasciatus* respectively (Table 1). This quality score improvement and reduced fragmentation supports use of the scaffolding approach, producing a higher-quality genome for downstream analyses.

**Table 1.** Comparison of genome quality metrics for Bermuda skink *Plestiodon longirostris* genome assemblies: a *de novo* assembly of Oxford Nanopore Technology long-reads, and assemblies produced from scaffolding the *de novo* assembly against *Plestiodon fasciatus* and *Plestiodon gilberti* reference genomes.

| **Metric** | ***De novo assembly*** | **Reference-scaffolded assemblies** |  |
|---|---|---|---|
|  |  | **Reference:**<br>***Plestiodon fasciatus*** | **Reference:**<br>***Plestiodon gilberti*** |
| Total size | 1.496 Gb | 1.496 Gb | 1.496 Gb |
| No. scaffold contigs | 13,098 | 2,579 | 3,391 |
| Scaffold N50 | 334 Kb | 224 Mb | 176 Mb |
| Largest scaffold | 2 Mb | 295.7 Mb | 262.2 Mb |
| No. macrochromosomes (>20 Mb) | n/a | 13 | 13 |
| GC content | 45.68% | 45.68% | 45.68% |
| BUSCO completedness | 94.1% | 98.70% | 98.20% |
| BUSCO missing | 2.3% | 0.90% | 1.00% |

Our assembly size corresponds with the size range of 1.5 – 2 Gb reported for other skink species (Hanrahan et al. 2025; Hoffman et al. 2026; Richmond et al. 2026). The assembly comprises 13 macrochromosomes, also within the expected range for a skink species of 11-16 (Deweese and Wright 1970; Giovannotti et al. 2010). This also matches the chromosome numbers of the reference genomes used (Hoffman et al. 2026; Richmond et al. 2026), though this may be due to bias introduced by the assembler assuming the karyotype is the same as the reference used, and karyotyping may be worth exploring further with this species. The remaining 2,565 unplaced contigs likely represent microchromosomal and repetitive sequences. We were unable to resolve microchromosomal content, which would require Hi-C proximity ligation data, a possible future study.

### Genomic and Transposable Element Landscapes

Using LiftOff, 94.8% (30,813 of 32,520) of gene models were mapped from *Plestiodon fasciatus* and 97.8% (18,408 of 18,814) from *P. gilberti.* Total repeat content was 48.37% of the genome, with 33.67% of the genome comprising unclassified TEs when scaffolded against both reference genomes. These values are comparable to other large squamate genomes, and the percentage of unclassified TEs is typical of non-model squamate species (Pasquesi et al. 2018). We found that LINEs made up 11.6% of our TEs and SINEs made up 0.7% of our TEs (Table 2). These values were the same against both scaffold assemblies, even with the total number of genes transferred, transfer efficiency, and unmapped genes varying slightly.

**Table 2.** Genome and Transposable Element landscape values for Bermuda skink *Plestiodon longirostris* Oxford Nanopore Technology long-read genomic data scaffolded against chromosome-level, annotated *P. fasciatus* and *P. gilberti* genomes.

| **Metric** | **Species used in reference-guided annotation** |  |
|---|---|---|
|  | ***Plestiodon fasciatus*** | ***Plestiodon gilberti*** |
|---|---|---|
| Internal stop codons | 9.00% | 9.00% |
| Total repeat content | 48.37% | 48.37% |
| Unclassified repeats | 33.79% | 33.81% |
| Retroelements | 12.26% | 12.25% |
| LTR elements | 1.98% | 1.98% |
| DNA transposons | 0.28% | 0.28% |
| Genes transferred | 30,813 | 18,408 |
| Reference gene count | 32,520 | 18,814 |
| Transfer efficiency | 94.80% | 97.80% |
| Unmapped genes | 5.20% | 2.20% |

While the proportion of internal stop codons in the assembly was 9.1%, comparable to the 9% of the *P. fasciatus* and *P. gilberti* genomes (Table 2; Hoffman et al. 2026; Richmond et al. 2026) and is not unusual in a Nanopore long-read assembly due to the elevated error rate in Nanopore reads. Moreover, we mitigated this using Pilon short-read polishing, and our BUSCO completeness and gene annotation completeness remained high despite it. Additionally, we had 5.2% unmapped genes from *P. fasciatus* through Liftoff, which could be due to annotation artefacts, or could be genuine gene absence or divergence, given their 11-19 million year divergence (Brandley et al. 2010). While our unmapped gene percentage for the *P. gilberti* assembly was lower, at 2.2%, this is possibly due to the fact there were fewer reference genes to transfer.

We found an increase in gene density from large macrochromosomes to smaller chromosomes, increasing from approximately 15–25 genes/Mb in chromosomes 1-6 to approximately 95 genes/Mb in chromosomes 7-13 (Figure 1). This inverse relationship between chromosome size and gene density is a conserved pattern typically seen in vertebrates (Pinto et al. 2023), consistent with an evolutionary origin of small macrochromosomes from ancestral microchromosomes (Waters et al. 2021). We also found an inverse relationship between gene density and repeat content, with gene-rich windows being repeat-poor (Figure 1). This is particularly notable in the smaller chromosomes 7-13 and supports their origination from microchromosomes.

**Figure 1.**
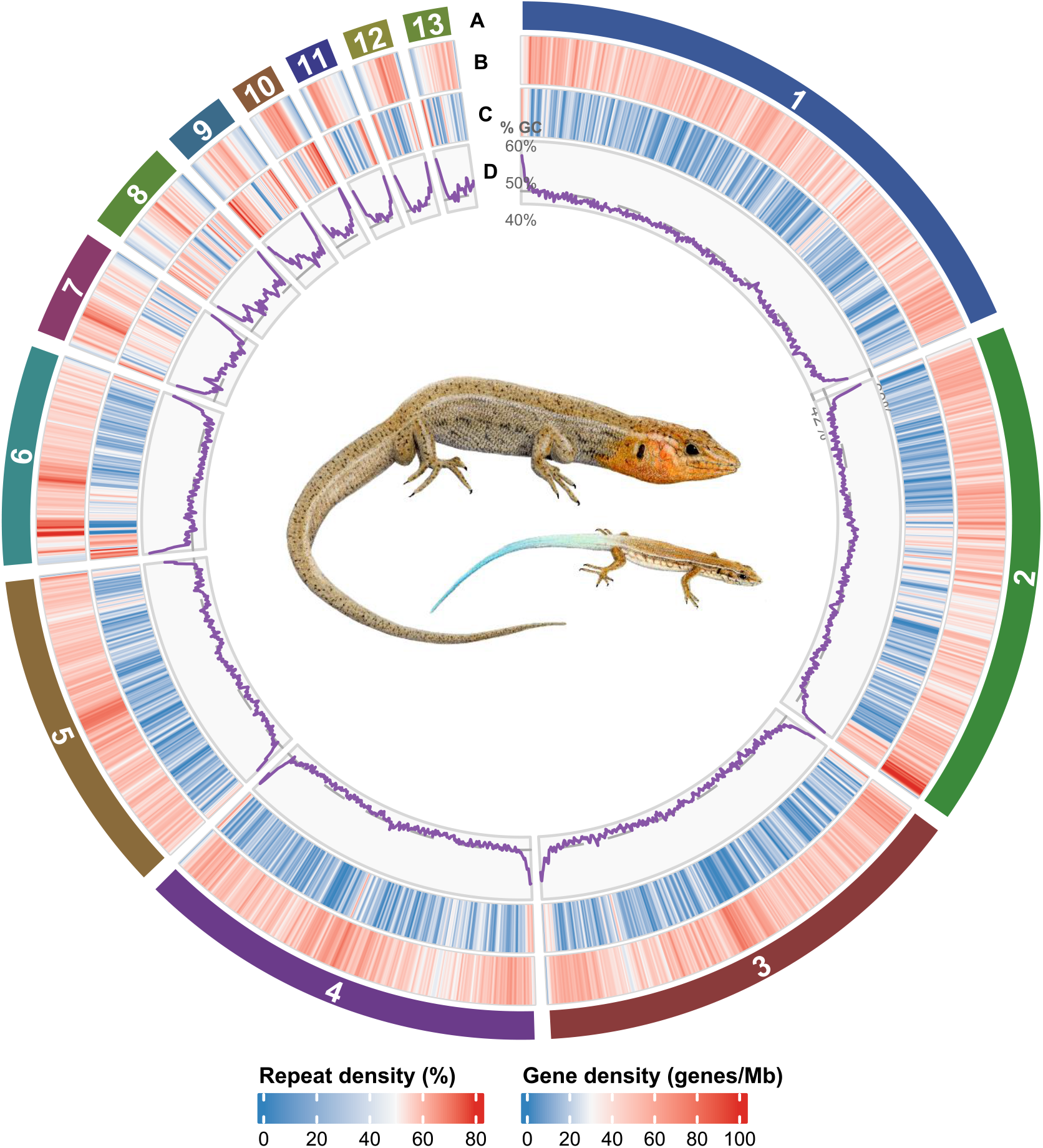
Ideogram for the Bermuda skink *Plestiodon longirostris*, with chromosome number and relative size (A), heatmaps of gene density (B) and repeat elements (C) in 50 kb windows, with darker colours for higher density, and a line chart of GC content across the genome (D). Circos was used to generate the plot (v0.69.6; Krzywinski et al., 2009). Scientific illustration of the adult (top) and juvenile (bottom) Bermuda skink by Dr. Guy Freeman. (Circos plot) Circular genome plot of the Bermuda skink (*Plestiodon longirostris*) showing 13 chromosomes arranged around a ring, decreasing in size from chromosome 1 to 13. Inner tracks show heatmaps of gene density and repeat density in 50 kb windows, colored from blue (low) through white to red (high). The innermost track is a line chart of GC content, ranging from about 40% to 60%. GC content is generally higher towards chromosome ends and on the smaller chromosomes.

The Kimura 2-parameter divergence landscape for the TEs in the genome peaked between 20-25%, suggesting the majority of TEs are ancient insertions, indicating no recent major TE bursts (Figure 2). This peak is consistent with expectations for a small island endemic, resulting from reduced effective population size limiting purifying selection against TE insertion, leading to TE persistence rather than purging of old copies.

**Figure 2.**
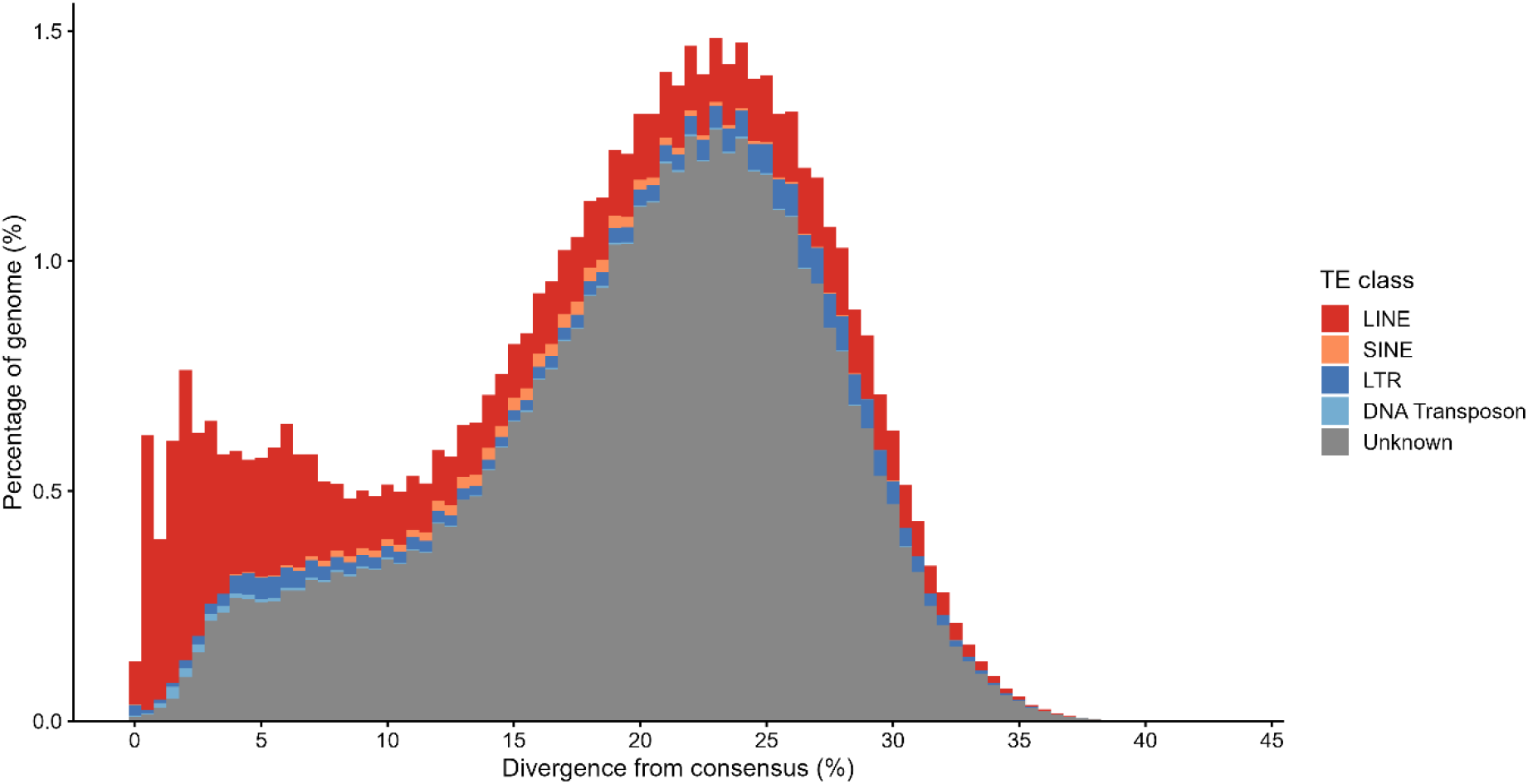
Transposable element (TE) landscape of Bermuda skink *Plestiodon longirostris,* based on Kimura 2-parameter (K2P) divergence by TE class. K2P divergence % shows the substitution distance between each TE copy and its consensus sequence, as a proxy for relative age, with lower divergence indicating younger elements. The y-axis shows the proportion of the genome occupied by each TE class at each divergence level. (TE landscape) Stacked bar chart of transposable element content against divergence from consensus (0–45%), colored by TE class: LINE, SINE, LTR, DNA transposon and unknown. The distribution has two peaks. A large, older peak around 20–25% divergence reaches about 1.5% of the genome and is mostly unclassified elements. A smaller, recent peak at 0–10% divergence is dominated by LINEs, indicating recent LINE activity.

However, we found increased LINE activity between 0-5% divergence, suggesting recent or ongoing LINE transposition activity, despite the TE landscape being predominantly ancient (Pasquesi et al. 2018). These ongoing LINE insertions could disrupt gene function and introduce new regulatory elements (Schrader and Schmitz 2019; Catlin and Josephs 2022), and in a Critically Endangered species with small effective population size where the forces of genetic drift are strong, could result in increased genetic load through accumulation of deleterious mutations (Pasquesi et al. 2018; Lynch and Conery 2003). However, compared to *P. fasciatus* (Hoffman et al. 2026), this shoulder is less pronounced, indicative of its isolation and smaller effective population size suppressing recent transposition.

Our TE landscape is predominantly ancient, and TE activity is not elevated. TE bursts have been associated with environmental stress, colonization of new habitats and population bottlenecks (Stapley et al. 2015; Schrader and Schmitz 2019), so the absence of recent activity suggests the Bermuda skink genome has not undergone such episodes in its recent evolutionary history. However, as previously mentioned, there may be ongoing gene function disruption, evidenced by current LINE transposition activity (0-5% divergence). Our high unclassified TE content (33.67%) (i.e. novel TEs not found in closest relatives) suggests a high proportion are lineage-specific or sufficiently diverged to be unclassified, common in understudied squamata lineages (Pinto et al. 2023; Pasquesi et al. 2018). Our use of *de novo* repeat modelling and generating a library through RepeatModeler provides a valuable resource for identifying these unique elements to advance understanding of squamate evolutionary history.

### Synteny Analysis

We found strong conservation of macrochromosomes and gene order across *Plestiodon* species, particularly between the Bermuda skink*, P. gilberti,* and *P. fasciatus* through riparian synteny analysis (Figure 3). We also found progressive chromosomal rearrangement as phylogenetic distance increased, most extensively in comparison with *Hemicordylus capensis* (∼156 Ma divergence). As macrochromosome size decreased, rearrangement increased across all comparisons, indicating the increased recombination rates and structural dynamism reported in microchromosome-derived elements in other squamate genomes (Hoffman et al. 2026; Richmond et al. 2026). Our synteny analysis also determined that chromosome 5 of the Bermuda skink is the putative X chromosome based on the *B. duperreyi* confirmed X chromosome (CM084881.2), consistent with *P. gilberti* and *P. fasciatus* (Hoffman et al. 2026; Richmond et al. 2026).

**Figure 3.**
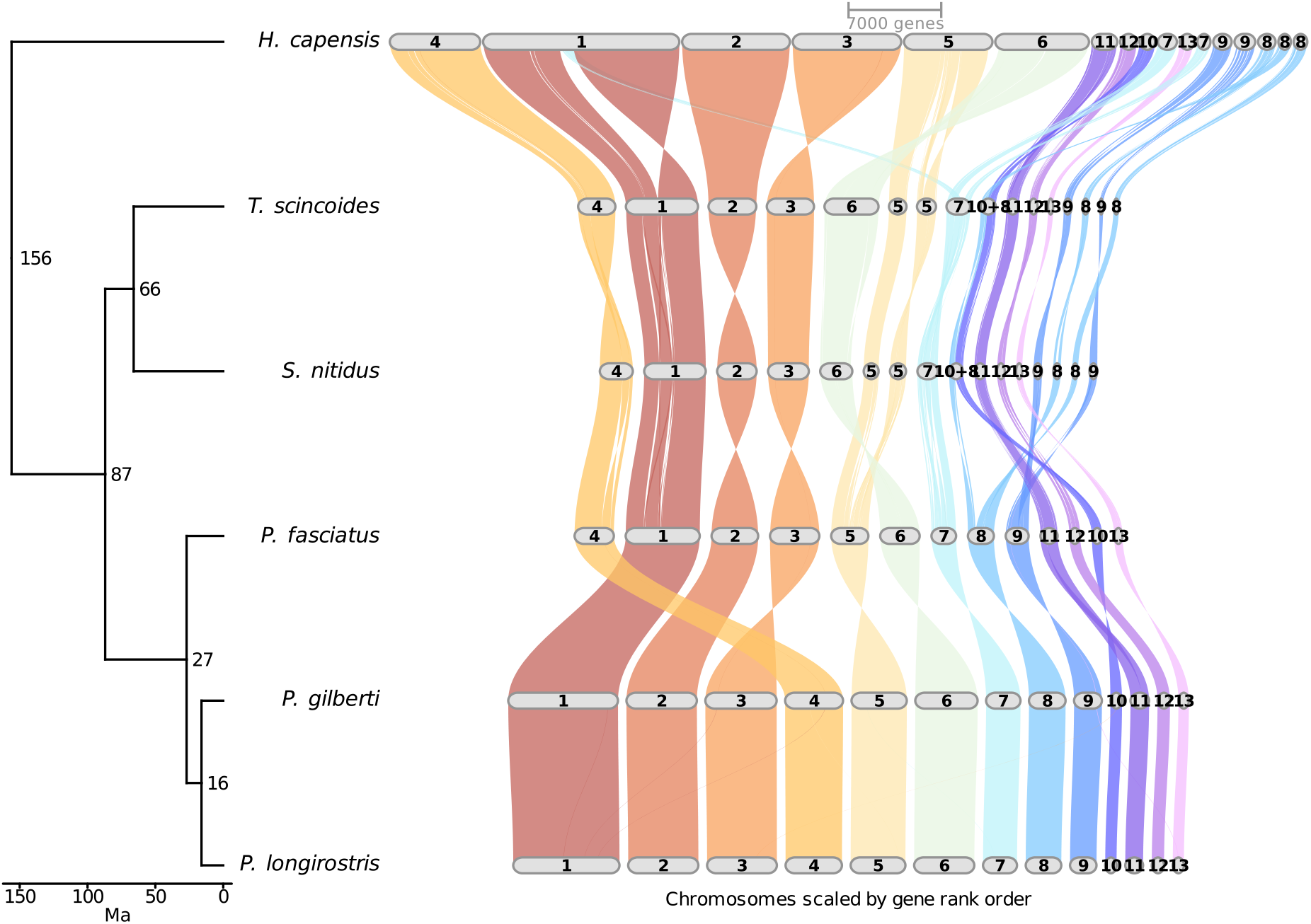
Riparian synteny plot across six skink species annotated genomes: Bermuda skink Plestiodon longirostris, P. gilberti, P. fasciatus, Tiliqua scincoides, Spondylurus nitidus and Hemicordylus capensis. (Synteny plot) Synteny plot linking the chromosomes of six species, with a time-calibrated phylogeny on the left. Estimated divergence times are 156, 87, 66, 27 and 16 million years ago, with the Bermuda skink and *Plestiodon gilberti* most closely related. Colored ribbons connect matching chromosomal regions between neighboring genomes. Large chromosomes are largely conserved across species, while the smaller chromosomes show more rearrangement, especially compared with H. capensis.

### Mitochondrial Genome

We assembled the mitochondrial genome of the reference individual from 24,400 Nanopore reads, with a mean depth of 2,748× and 99.8% of positions covered. The final polished assembly was 17,415 bp long, consistent with the typical vertebrate mitochondrial genome size range of 16–18 kb, with a GC content of 43.1%. Validation by remapping the reads identified six positions in the control region where the majority of reads disagreed with the polished consensus; after correcting these, remapping gave no variants at ≥50× depth. MITOS2 annotation identified the complete vertebrate mitochondrial gene complement (Figure 4): 13 protein-coding genes (ND1, ND2, COX1, COX2, ATP8, ATP6, COX3, ND3, ND4L, ND4, ND5, ND6, CYTB), 22 transfer RNA genes, two ribosomal RNA genes (12S and 16S) and the control region (D-loop). Gene order and strand orientation were consistent with the ancestral vertebrate arrangement, with 12 protein-coding genes on the heavy strand and ND6 on the light strand.

**Figure 4.**
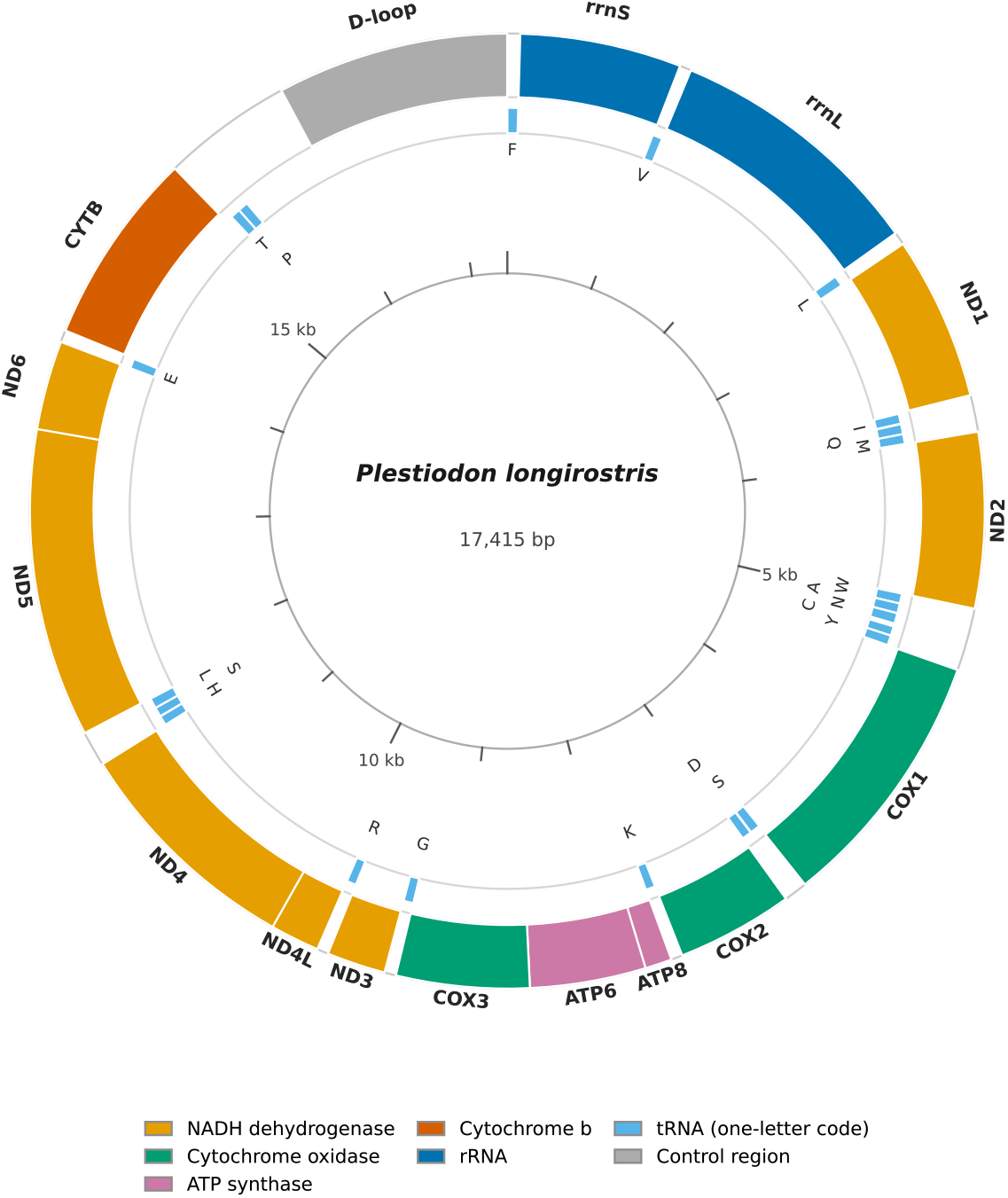
Annotated mitochondrial genome for the Bermuda skink *Plestiodon longirostris,* assembled from Oxford Nanopore reads. (Mitochondrial genome) Circular map of the 17,415 bp mitochondrial genome of the Bermuda skink. It shows 13 protein-coding genes (NADH dehydrogenase, cytochrome oxidase, ATP synthase and cytochrome b subunits), two rRNA genes, 22 tRNA genes labelled by one-letter code, and the D-loop control region. Clockwise from the top, the gene order is rrnS, rrnL, ND1, ND2, COX1, COX2, ATP8, ATP6, COX3, ND3, ND4L, ND4, ND5, ND6, CYTB and the D-loop.

## Conclusion

This chromosome-level annotated genome assembly represents a significant contribution to Bermuda skink genomics, and will be important in supporting conservation efforts for this Critically Endangered, paleoendemic species. It could support demographic reconstruction, as well as population genomic resequencing studies, allowing investigation into deleterious mutation load, inbreeding, dispersal and gene flow dynamics, genetic diversity, adaptive potential and patterns of selection. This would improve our understanding of resilience and vulnerability to genetic and demographic threats in the species, as well as enabling the development of more targeted and effective conservation management strategies. Furthermore, this genome represents the last remaining species of an evolutionary lineage, and can be used in comparative genomic studies across the Scincidae family and wider reptile group to investigate broader evolutionary patterns and mechanisms.

## Data Availability

The Bermuda skink *Plestiodon longirostris* genome sequence is available upon request and is available through NCBI Bioproject: PRJNA1479328. Sequences used to assemble the genome are available through the SRA under the same accession. Short-read sequences used to polish the genome are available through NCBI Bioproject: PRJNA1558259. All code used to generate data is available at <u>owengw/Plong-genome-pipeline</u> and <u>owengw/mito_pipeline</u>. All other genomes used are referenced and where an accession number is provided are publicly available on NCBI.

## Acknowledgements

Thanks to Hans Sebastian Manrique Castaneda, Natacha Smith, Olivia Lee, Joshua Wright, John Weale, Heather and Eva Bottelli, Daniel Dickinson and Jessica Young for their assistance in the field. Thanks to Jeremy Madeiros and Peter Drew for providing access to and support in Protected Areas in Castle Harbour. Claude Sonnet 4.6 was used to optimize and annotate autosomal and mitochondrial genome scripts with scripts I drafted and provided, which output was then reviewed by Owen Greenwood.

## Funding

Thanks to Oxford Nanopore Technology’s Grant Programme, which enabled the generation of the long-read sequence data through BioQuest. NEOF and the Centre for Genomics Research, University of Liverpool, which funded the sequencing of the short-read sequence data. This study was supported and funded by Manchester Metropolitan University, Chester Zoo, the Bermuda Zoological Society, The Genetics Society and the British Herpetological Society.

## Conflicts of interest

None of the authors have any conflicts of interest to declare.

## Author Contributions

Conceptualization, editing drafts of the manuscript: all authors; methodology design: OG, KB-B, HR, KJ-M and CW; data analysis: OG, MJ; writing original manuscript draft: OG; funding acquisition: OG, LW, CW and SG.

## Ethical Standards

All work handling and sampling Bermuda skinks was approved by the Manchester Metropolitan University Ethics panel (EthOS Project ID: 50781) and the Chester Zoo Ethical Review Committee.

